# Identification of subject-specific immunoglobulin alleles from expressed repertoire sequencing data

**DOI:** 10.1101/405704

**Authors:** Daniel Gadala-Maria, Moriah Gidoni, Susanna Marquez, Jason A. Vander Heiden, Justin T. Kos, Corey T. Watson, Kevin C. O’Connor, Gur Yaari, Steven H. Kleinstein

## Abstract

The adaptive immune receptor repertoire (AIRR) contains information on an individuals’ immune past, present and potential in the form of the evolving sequences that encode the B cell receptor (BCR) repertoire. AIRR sequencing (AIRR-seq) studies rely on databases of known BCR germline variable (V), diversity (D) and joining (J) genes to detect somatic mutations in AIRR-seq data via comparison to the best-aligning database alleles. However, it has been shown that these databases are far from complete, leading to systematic misidentification of mutated positions in subsets of sample sequences. We previously presented TIgGER, a computational method to identify subject-specific V gene genotypes, including the presence of novel V gene alleles, directly from AIRR-seq data. However, the original algorithm was unable to detect alleles that differed by more than 5 single nucleotide polymorphisms (SNPs) from a database allele. Here we present and apply an improved version of the TIgGER algorithm which can detect alleles that differ by any number of SNPs from the nearest database allele, and can construct subject-specific genotypes with minimal prior information. TIgGER predictions are validated both computationally (using a leave-one-out strategy) and experimentally (using genomic sequencing), resulting in the addition of three new immunoglobulin heavy chain V (IGHV) gene alleles to the IMGT repertoire. Finally, we develop a Bayesian strategy to provide a confidence estimate associated with genotype calls. All together, these methods allow for much higher accuracy in germline allele assignment, an essential step in AIRR-seq studies.

## INTRODUCTION

Affinity maturation, in which B cells expressing receptors with an improved ability to bind antigen are preferentially expanded, is a key component of the B cell-mediated adaptive immune response [1,2]. This selection process requires a diverse pool of B cell receptors (BCRs) which is generated both through V(D)J recombination (in which each B cell creates its own BCR by recombining variable (V), diversity (D) and joining (J) genes), and through the subsequent somatic hypermutation (SHM) of these sequences during T-dependent adaptive immune responses. SHM is an enzymatically-driven process that introduces mainly point substitutions into the BCR at a rate of about one mutation per 1,000 base-pairs per cell division [3,4]. Leveraging next-generation sequencing technologies to profile this adaptive immune receptor repertoire (AIRR) allows tens- to hundreds-of-millions of unique BCR sequences to be collected from a single subject and has become a prevalent method for studying aspects of the B cell-mediated immune response, including topics related to gene usage, mutation patterns, and clonality [5-9].

An accurate immunoglobulin (Ig) germline receptor database (IgGRdb) is a key part of the typical AIRR-seq data analysis pipeline [10]. Analysis generally begins with pre-processing tools specifically designed for BCR sequencing, such as pRESTO [11]. Following this, computational methods (e.g., IMGT/HighV-QUEST [12], IgBLAST [13], or iHMMune-Align [14]) are used to align sample sequences to the set of unmutated reference alleles from an IgGRdb, such as the one maintained by IMGT [3]. However, these IgGRdbs have been shown to be incomplete, and studies continue to discover new alleles [5-9]. Immunoglobulin (Ig) loci are rarely fully sequenced in a single subject due to the large locus size and similarity of genes confounding many modern high-throughput sequencing methods [7,15,16]. Thus, if a subject carries a novel allele, it can lead to incorrect interpretations of which positions have been mutated and can subsequently affect the reconstruction of clonal lineages. We previously created the TIgGER method, and an associated software package, to detect novel V gene alleles from AIRR-seq data, infer the genotype of a subject, and correct the initial allele assignments [8]. While the application of TIgGER to several subjects revealed a high prevalence of novel alleles, the design of the method limited its ability to detect novel alleles differing by more than five polymorphisms from a known IgGRdb allele, which we estimated would account for ~12% of novel alleles [8].

Here we present and apply improvements upon the original TIgGER method that allow for the detection of novel alleles that differ greatly from IgGRdb alleles as well as for the assignment of levels of confidence to each genotype call. This updated version of TIgGER (version XX or higher – during the review process the source code is available at the tip of the TIgGER bitbucket repository at https://bitbucket.org/kleinstein/tigger) is available for download as an R package from The Comprehensive R Archive Network (CRAN; http://cran.r-project.org), with additional documentation available at http://tigger.readthedocs.io. The input and output formats of TIgGER conform to the Change-O file standard [17], and thus the method can be used seamlessly as part of the Immcantation tool suite, which provides a start-to-finish analytical ecosystem for high-throughput AIRR-seq datasets (http://immcantation.org), including methods for pre-processing, population structure determination, and advanced repertoire analyses.

## RESULTS

### Detecting distant alleles using dynamic positioning of the “mutation window”

TIgGER detects novel alleles by analyzing the apparent mutation frequency pattern at each nucleotide position as a function of the sequence-wide mutation count. The input to the method consists of a set of rearranged BCR sequences from a single subject and those sequences’ alignments to IgGRdb alleles, such as the output of running IMGT/HighV-QUEST [4,18]. TIgGER searches for novel V gene alleles among the sequences that fall in a specified “mutation window” relative to each of the IgGRdb alleles. The mutation window of the original algorithm [8] had an upper bound of at most 10 sequence-wide mutations, while the lower bound was defined as *minimum(*L, 5), where L was the most frequent mutation count among sequences with at most 10 sequence-wide mutations. Positions were considered as potentially polymorphic if a linear fit predicted a mutation frequency (y value) above a threshold level of 0.125 at a mutation count (x value) of zero (i.e., the y-intercept). While this method had excellent sensitivity and specificity, the definition of the lower bound meant that TIgGER could only detect novel alleles that differed by at most 5 single nucleotide polymorphisms from some previously known IgGRdb allele. We hypothesized that by modifying the TIgGER algorithm to dynamically shift the mutation window to the most relevant region for discovery of the polymorphic position, it would be possible to detect novel V gene alleles that differed by any number of polymorphisms from the nearest IgGRdb allele.

The updated TIgGER algorithm described here defines the lower bound of the mutation window for each allele as the mutation count of the most frequent sequence assigned to that allele. The upper bound of the mutation window is always nine bases greater than the lower bound. Positions are analyzed within this window, and considered as potentially polymorphic if a linear fit predicts a mutation frequency (y value) above a threshold level of 0.125 at a mutation count (x value) one less than the start of the mutation window (see Methods for details). The behavior of the updated TIgGER algorithm (Figure 1, bottom row) is equivalent to the original TIgGER algorithm (Figure 1, top row) when analyzing sequences derived from a novel allele with a single nucleotide polymorphism (Figure 1, first column). The behavior of the two algorithms diverges slightly in cases where 2-to-5 polymorphisms are present in the novel allele (Figure 1, middle column), as the updated algorithm allows both the upper bound of the mutation window and the location where the mutation frequency threshold is evaluated to dynamically shift based on the start of the window. The greatest divergence is observed in detecting novel alleles with over 5 single nucleotide polymorphisms. In this case, the mutation window of the original algorithm ends before the window of the updated algorithm (Figure 1, right column). When confronted with such distant novel alleles, the linear fits of the polymorphic positions constructed by the original algorithm often failed to yield y-intercepts large enough to identify the positions as polymorphic, whereas the updated algorithm can identify all polymorphic positions.

**Figure 1:**
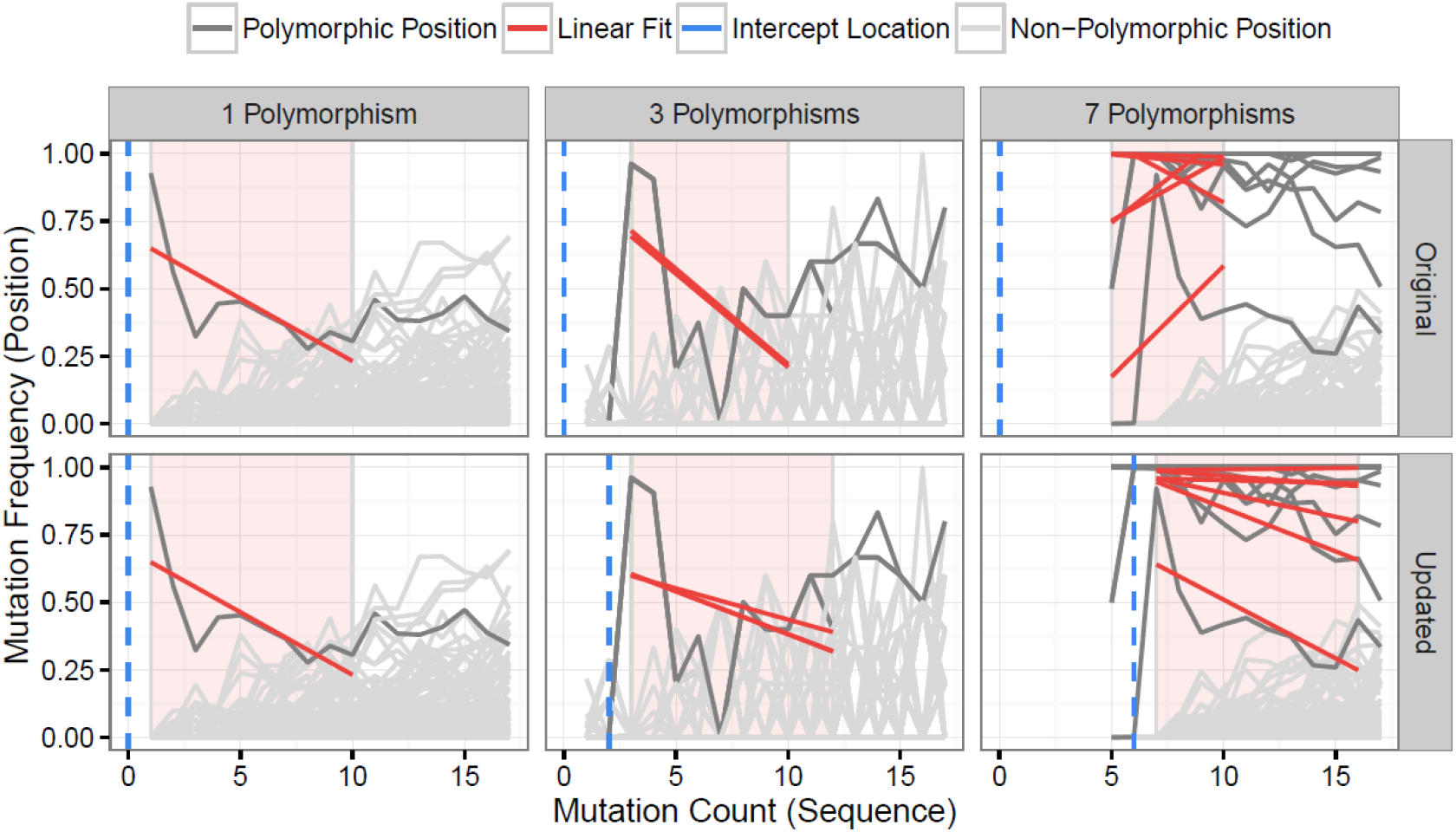
Distant V gene alleles can be detected by dynamic shifting of the mutation window. The original TIgGER algorithm (top row) and the updated method (bottom row) were applied to BCR sequences generated from two subjects, hu420143 and 420IV, as part of a vaccination time course study (16). In both cases, the mutation frequency (y-axis) at each nucleotide position (grey lines) was determined as a function of the sequence-wide mutation count (x-axis). For each position known to be polymorphic (dark grey lines) (12), linear fits (red lines) were constructed using the points within the mutation window (red shaded region). The linear fit was then used to estimate the mutation frequency at the intercept location (blue dotted line). Sequences that best aligned to IGHV1-2*02 from hu420143 were used to demonstrate the behavior when detecting a germline with a single nucleotide polymorphism (left column), while sequences that best aligned to IGHV3-43*01 from 420IV were used to demonstrate the behavior when detecting a germline with three polymorphisms (middle column), as novel alleles with that number of polymorphisms had been previously discovered in those subjects (12). Data to assess the behavior when detecting a novel allele with seven polymorphisms (right column) was simulated using sequences from hu420143 that best aligned to IGHV1-2*02 by artificially adding six base changes to the germline sequence used for alignment, as no novel allele with more than five polymorphisms had been discovered. In all cases, only sequences from pre-vaccination time points were used from these individuals.

To test the performance of the updated TIgGER method, we simulated data in which novel alleles differed by *n* SNPs from the nearest IgGRdb allele by randomly changing *n* nucleotides in the IgGRdb alleles utilized by TIgGER (i.e., by removing the true allele from the IgGRdb and replacing it with a distant one). Using AIRR-seq data from subject PGP1 described in our previous study [19], the 38 IGHV alleles assigned to at least 500 unique BCR sequences were each tested for every value of *n* from 1 to 30. This process was repeated 100 times per value of *n* random single nucleotide polymorphisms, to ensure a diversity of polymorphic positions and base changes would be tested. The fraction of times the original germline sequence was recovered was determined as a function of *n* and averaged across all germline alleles tested. The updated version of TIgGER had 100% sensitivity in the range of 1 ≤ *n* ≤ 5, and was also able to detect novel alleles with high sensitivity (over 99%) for all values of *n* tested (Figure 2). Additionally, only the removed germline alleles were discovered by the algorithm; no false positive sequences were predicted. Thus, TIgGER can detect novel V gene alleles that are far from any known IgGRdb allele with high sensitivity and specificity.

**Figure 2:**
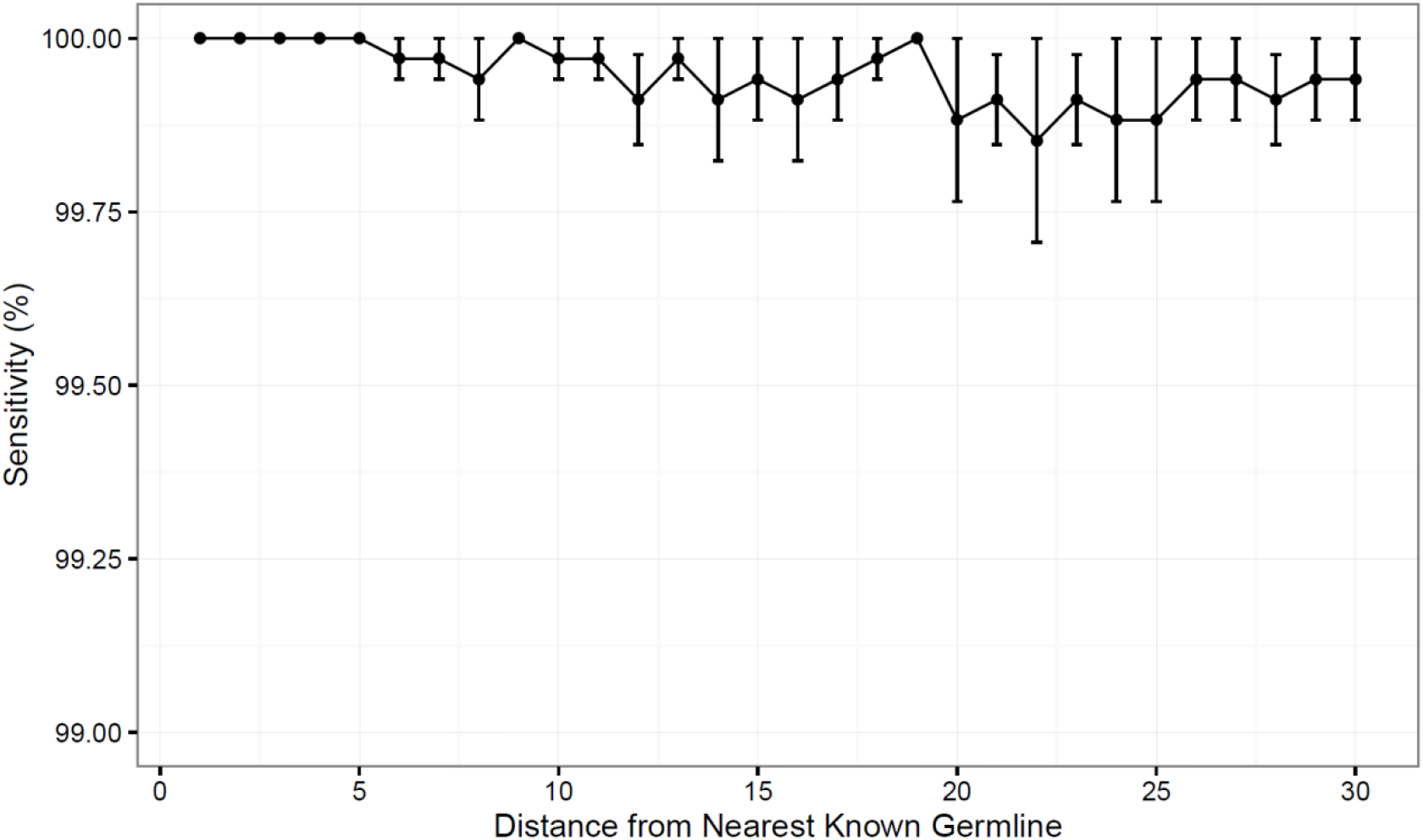
The updated TIgGER method detects distant alleles with high sensitivity. Detection of novel V gene alleles differing from IgGRdb alleles by *n* polymorphisms was simulated by extracting experimental sequences best aligning to a single IgGRdb allele in a single subject, then inserting into the IgGRdb allele *n* polymorphisms *in silico* and proving only the modified IgGRdb allele to TIgGER. Each sensitivity measurement at distance (x-axis) included modification of all IgGRdb alleles best-aligning to at least 500 sequences in subject PGP1. The variance in sensitivity was estimated by repeating this procedure for 100 randomly-modified IgGRdb alleles and the mean sensitivity as a function of *n* was determined for 1 ≤ *n* ≤ 30. Error bars represent the standard error of the mean.

Based on the distances between alleles in the IMGT IgGRdb, we previously estimated that ~12% of novel alleles should differ by more than five polymorphisms from the nearest IgGRdb allele. To search for such distant novel alleles, the updated version of TIgGER was applied to AIRR-seq data from the seven individuals described in our previous study [8], including three subjects receiving influenza vaccination [19] and four subjects with multiple sclerosis [20,21]. However, this yielded the same alleles previously reported, with the most-distant novel alleles differing from the nearest IgGRdb allele by at most three polymorphisms (Table S1). We next applied the updated TIgGER algorithm to 24 additional individuals. This included published AIRR-seq data from five pairs of monozygotic twins (10), 10 subjects with myasthenia gravis and 4 subjects that served as healthy controls [22]. Considering all 31 individuals, TIgGER identified a total of 28 novel alleles that were part of the genotype inferred for one or more of the individuals (Figure 3 and Table S1). All of the novel alleles differed from IgGRdb alleles by at most three single nucleotide polymorphisms. Thus, while it was demonstrated on synthetic data that the updated version of TIgGER has the potential to detect alleles that differ greatly from known IgGRdb alleles, none of the novel alleles discovered in the repertoires of 26 genetically distinct individuals (monozygotic twins are considered genetically indistinguishable) differed by more than three polymorphisms.

**Figure 3:**
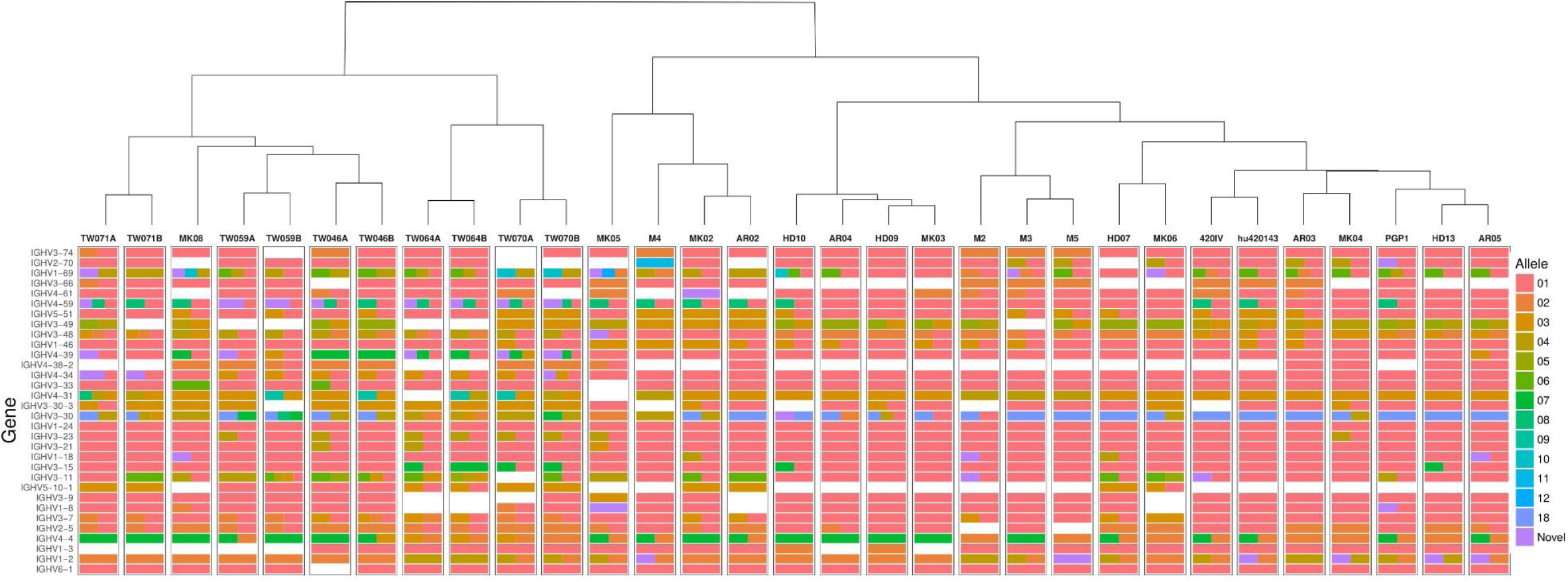
TIgGER identifies high diversity in the inferred IGHV genotypes of 26 genetically distinct subjects, and high similarity among 5 monozygotic twin pairs. IGHV genotypes of 31 subjects (columns) were inferred using the frequency method from the updated TIgGER algorithm, under the default settings. Monozygotic twin pairs are indicated as TW***A and TW***B; multiple sclerosis subjects are denoted as M*; influenza vaccination time course subjects are denoted PGP1, 420IV, and hu420143; myasthenis gravis subjects are indicated by AR** and MK** with the associated healthy controls indicated by HD**. Each row represents an IGHV gene, and the color(s) in each column represent(s) the allele(s) present in that subject using the IMGT designation shown in the color bar. Note that the size of the bars does not represent relative allele frequency. Clustering of the genotypes was performed using Ward’s method.

### Experimental validation of novel IGHV gene alleles predicted by TIgGER

The application of TIgGER to AIRR-seq data from 26 genetically distinct individuals identified 28 novel IGHV gene alleles (Figure 3 and Table S1). We selected four of these novel alleles that were each predicted by TIgGER in multiple individuals for experimental validation: *IGHV1-2*02_T163C*, *IGHV1-8*02_G234T*, *IGHV3-20*01_C307T* and *IGHV1-69*06_C191T*. Three of these alleles were also predicted independently by other groups. *IGHV1-2*02_T163C* was identified in [5,9], *IGHV1-8*02_G234T* was identified in [9] and *IGHV3-20*01_C307T* was identified in [23]. *IGHV1-69*06_C191T* has not been previously reported.

To validate the TIgGER predictions, we cloned and sequenced the relevant gene locus directly from genomic DNA. For each allele, we chose one of the subjects where it was predicted for validation: MK04, MK05, MK05 and MK06 for the alleles of *IGHV1-2*, *IGHV1-8*, *IGHV3-20* and *IGHV1-69*, respectively. PCR primers were designed to fully amplify the exons and introns of each target IGHV gene locus (*IGHV1-2*, *IGHV1-8*, *IGHV3-20* and *IGHV1-69*) from genomic DNA; sequences for each primer set are provided in Table S2. PCR amplicons for each gene were then generated individually from the genomic DNA samples of the donor where they were predicted to be present, and subsequently cloned. DNA was isolated from 4-15 clones per gene target, and Sanger sequenced from both ends. These sequences were compared directly to the allele sequences inferred by TIgGER from the same donor to assess the degree of concordance. In all cases (4/4), genomic DNA sequencing provided validation of the putative IGHV polymorphisms inferred by TIgGER from the AIRR-Seq data suggesting that TIgGER has high specificity for identifying new IGHV alleles.

Single representative clones for each genomic sequence validating the TIgGER predictions were submitted to GenBank and have been assigned the following accession numbers: MH267285 (*IGHV1-2*02_T163T*), MH267286 (*IGHV1-8*02_G234T*), MH332884 (*IGHV3-20*01_C307T*) and MH359407 (*IGHV1-69*06_C191T*). These predicted alleles were also submitted to IMGT for inclusion in their IgGRdb. Three of these alleles were accepted for inclusion in the IMGT IgGRdb as novel alleles, and have been assigned the following allele names: *IGHV1-2*06* (MH267285), *IGHV3-20*03* (MH332884) and *IGHV1-69*17* (MH359407). The fourth allele that we experimentally validated (*IGHV1-8*02_G234T*) was added to the IMGT IgGRdb as *IGHV1-8*03* during the course of this study, and was thus no longer considered novel. Along with *IGHV1-8*03*, several other alleles identical to TIgGER predictions were added to IMGT during this study: *IGHV1-18*01_T111C* as *IGHV1-18*04*, *IGHV2-70*01_T164G* as *IGHV2-70*15*, *IGHV3- 64*05_A210C_G265C* as *IGHV3-64D*06* and *IGHV3-9*01_C296T* as *IGHV3-9*03*. Overall, eight of the 28 novel IGHV genes predicted by TIgGER in 26 genetically distinct individuals are now part of the IMGT IgGRdb, including three novel IGHV alleles that directly resulted from this study.

### Inference of IGHV genes starting from a sparse IgGRdb

TIgGER relies on the ability to make initial assignments of BCR sequences to alleles from an IgGRdb. However, such IgGRdbs may be sparse or non-existent for certain species; IMGT/GENE-DB has only a single IgGRdb IGHV allele for most genes in mouse, and only a single allele for all genes in rat and rhesus macaque. Nevertheless, IGHV variation was observed in all of these species (for example, Mouse [24,25], Rat [26], Macaque [27,28]). In principle, the deep coverage of repertoire sequencing data could obviate the need for IgGRdbs by inferring the set of alleles for each subject based solely on the observed set of rearranged sequences. Here we consider whether a very sparse IgGRdb may be sufficient to discover the IGHV gene alleles of a subject’s IGHV genotype. This is theoretically possible given the ability of the updated TIgGER algorithm to detect alleles that differ greatly from the nearest known IgGRdb allele.

To evaluate the ability of TIgGER to identify the set of alleles carried by an individual when starting from a sparse IgGRdb, we simulated the extreme case of each gene family containing only a single allele in the IgGRdb. The performance was evaluated on published sequencing data from three subjects (PGP1, hu420143, and 420IV; see Methods). For each subject, the IgGRdb was defined by the single alleles from each IGHV family that were most frequently assigned by IMGT/HighV-QUEST. All sequences initially assigned to any allele in that family were then reassigned to that single IgGRdb allele. The set of IGHV genes carried by each individual was then identified by iterative applications of TIgGER. After each application of TIgGER, the set of novel alleles discovered by running the algorithm was added to the IgGRdb to be used for subsequent iterations, and sequences were reassigned to their most similar IgGRdb allele (measured by Hamming distance). The process was repeated until no new allele assignments were made (at most five iterations in these studies). The final set of alleles of each IGHV family discovered by this method was compared to the result obtained when running the TIgGER algorithm followed by genotype inference using the original IMGT/HighV-QUEST allele assignments and full IgGRdb (Figure 4).

**Figure 4:**
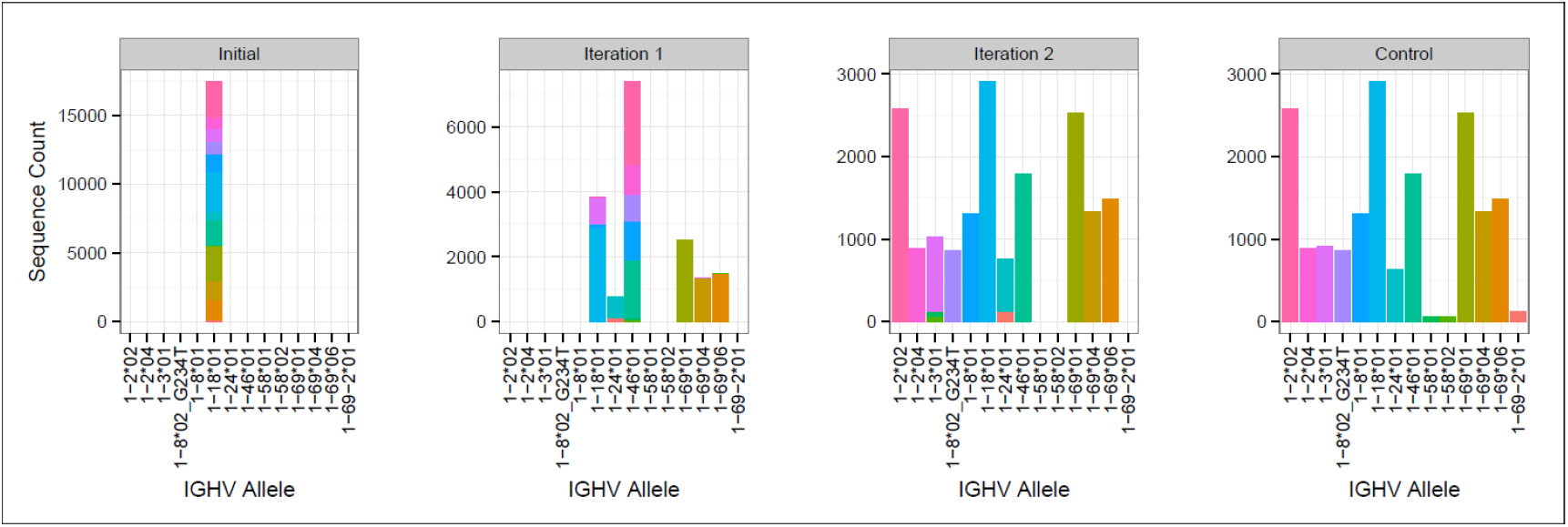
Iterative application of TIgGER detects most V gene alleles beginning from a sparse IgGRdb containing only one allele per IGHV family. Sequences from subject PGP1 that best aligned to any IgGRdb allele from the IGHV1 family were all reassigned to the most common allele, IGHV1-18*01 (first panel). The updated TIgGER algorithm was used to detect “novel” alleles that were not in the IgGRdb (initially consisting only of IGHV1-18*01), and then sample sequences were reassigned to the nearest previously-known or newly-discovered allele (second panel). After a second iteration (third panel), the sequence assignments closely matched the control case resulting from providing TIgGER with all known IGHV IgGRdb alleles (fourth panel). Bar colors indicate the allele assignments given to sample sequences in the control case.

**Table 1:** Performance of TIgGER in detecting the set of V gene alleles comprising each IGHV family starting from a sparse IgGRdb. As depicted in Figure 3, TIgGER was run iteratively to detect the set of alleles carried by three subjects. In each case, the algorithm was provided an initial IgGRdb consisting of only the single most-commonly observed allele for each IGHV family. Performance was assessed by comparing the final number of alleles per family discovered by this iterative method to the number of alleles per family resulting from running the TIgGER algorithm when provided with a complete list of IgGRdb alleles.

| Subject | IGHV Family | Alleles Discovered / Alleles Present (%) |
| --- | --- | --- |
| PGP1 | 1 | 11/14 (79%) |
| PGP1 | 2 | 6/6 (100%) |
| PGP1 | 3 | 14/29 (48%) |
| PGP1 | 4 | 10/13 (77%) |
| PGP1 | 5 | 1/2 (50%) |
| PGP1 | 6 | 1/1 (100%) |
| PGP1 | 7 | 1/2 (50%) |
| hu420143 | 1 | 8/12 (67%) |
| hu420143 | 2 | 5/5 (100%) |
| hu420143 | 3 | 16/22 (73%) |
| hu420143 | 4 | 9/11 (82%) |
| hu420143 | 5 | 1/1 (100%) |
| hu420143 | 6 | 1/1 (100%) |
| hu420143 | 7 | 1/1 (100%) |
| 420IV | 1 | 12/12 (100%) |
| 420IV | 2 | 5/5 (100%) |
| 420IV | 3 | 24/27 (89%) |
| 420IV | 4 | 9/9 (100%) |
| 420IV | 5 | 3/3 (100%) |
| 420IV | 6 | 1/1 (100%) |
| 420IV | 7 | 1/1 (100%) |
| **TOTAL** |  | **140/178 (79%)** |

**The updated TIgGER algorithm discovered an average of 79% of the alleles in each of the three subjects’ IGHV families when starting with a single IgGRdb allele per family (Table 1** Table 1). To understand how TIgGER achieves this performance, consider sequences from the IGHV1 family in subject PGP1. In this case, the first application of TIgGER was able to identify five of the correct novel alleles and reassign the sequences to the better allele (Figure 4, first and second panels). This success was due to the fact that the mutation ranges of interest (i.e., the mutation windows described in Figure 1) differed for many of the novel alleles. We expect this will generally be true, and since the number of positions differentiating different novel alleles from a shared most-similar IgGRdb allele varies, relevant mutation windows of alleles to be discovered are unlikely to overlap and result in a dilution of signal. Nevertheless, a single run of TIgGER was not able to detect all of the IGHV alleles. TIgGER was then run a second time using the new IgGRdb and assignments determined from the first run, leading to the identification of five additional novel alleles. This second iteration discovered less-used alleles, as the initial group of sequences assigned to the starting allele was broken into smaller subgroups (Figure 4, third panel). Three low-frequency alleles from two genes present when running TIgGER with access to the full IgGRdb (Figure 4, fourth panel) remained undiscovered after repeated iterations. Overall, these results demonstrate that TIgGER can be run iteratively to discover a large fraction of the IGHV alleles carried by an individual, even when there is very little prior knowledge of the set of alleles in the population.

### Bayesian inference of BCR genotypes can differentiate subjects

Given the diverse nature of the IGHV locus [7], we expected that genotypes inferred by TIgGER would vary across unrelated subjects, but should be the same within the five pairs of monozygotic twins. While the genotypes that were constructed for the individuals in this study were observed to be unique across subjects, the inferred genotypes of the monozygotic twin pairs were similar but not identical (Figure 3). Due to the relatively small number of sequences, not all novel alleles discovered in one twin were also discovered in the other. However, for the majority of genes, TIgGER assigned the same genotype alleles to each twin. Additionally, hierarchical clustering (using Ward’s method) of the genotypes properly grouped pairs of twins and excluded the genotypes of the other subjects (Figure 3, top).

In order to quantify our confidence in the assignment of genotypes, a Bayesian approach to genotyping was developed. This method analyzes the posterior probabilities of possible allele distributions, considering up to four distinct alleles per V. The posterior probabilities for these four possible models are compared and a Bayes factor is calculated for the two most probable models (see Methods). This Bayes factor reflects our confidence in the genotyping call of the method, and different models (i.e., different combinations of alleles) can be compared in a quantitative way. In the current implementation of the Bayesian approach, up to four alleles are considered (14), allowing for the possibility of a gene duplication with both loci being heterozygous (see Methods). This Bayesian method was applied separately to ten independent samples from subjects PGP1, hu420143 and 420IV (corresponding to 10 different time-points pre- and post-influenza vaccination) to test if we could confidently group samples from the same subject. The similarity of these personalized genotypes (for each combination of subject and time point) was estimated by determining the Jaccard distance metric for each gene. These individual gene distances were combined by calculating a weighted average of them using the Bayes factors as weights (see Methods). Using this distance metric, all samples from the three subjects could be differentiated with perfect accuracy, as the maximal weighted Jaccard distance of samples coming from the same subject was lower than the distance between samples coming from different subjects (Figure 5). Similar high classification accuracy was found for a wide range of model parameters showing the robustness of this approach. Overall, this Bayesian approach enables us to relax the strict cutoff criterion used by TIgGER in the previous sections (wherein the minimum number of alleles explaining 88% (7/8) of apparently-unmutated sequences are included in the genotype) to decide whether an allele should be included in an individual’s genotype or not.

**Figure 5:**
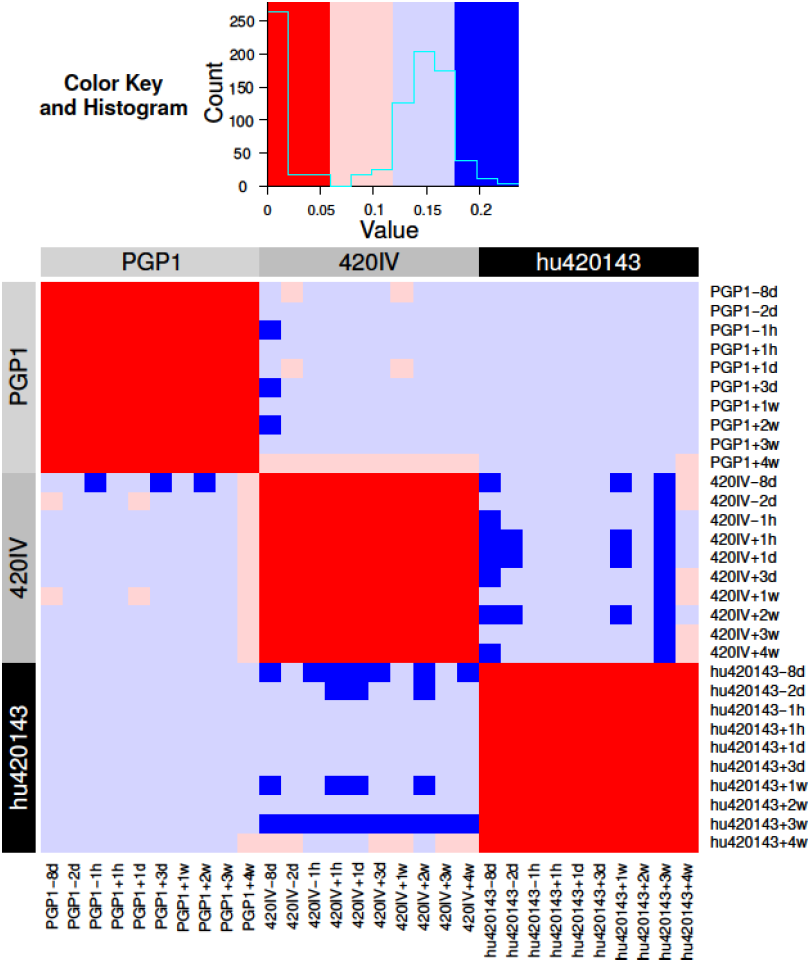
Bayesian inference of the subject-specific IGHV genotype identifies the same subject across independent samples in an influenza vaccination time course. BCR repertoire sequencing was carried out from a total of ten blood samples taken before and after influenza vaccination of three subjects (PGP1, 420IV, and hu420143) as part of a previous study (16). The Bayesian model was applied to the data from each of ten time points from each individual separately to determine a subject-specific IGHV genotype. The distance (colors) between each pair of inferred genotypes (rows/columns; numbered 1 – 30, labeled by color according to subject) is based on the Jaccard distance of the alleles of each gene (see Methods for details).

## METHODS

### Sample preparation, sequencing and processing of influenza vaccination data

Data from subjects PGP1, hu420143, and 420IV result from previously published BCR sequencing from blood samples taken at ten times relative to the administration of an influenza vaccine: -8 days, -2 days, -1 hour, +1 hour, +1 day, +3 days, +7 days, +14 days, +21 days, and +28 days. Peripheral blood was collected under the approval of the Personal Genome Project. Samples were prepared, sequenced and processed as described [19]. Briefly, V_H_ mRNA was selectively amplified by PCR using IGHV and IGHC region specific primers followed by sequencing on the Roche 454 platform. Sequence data were quality controlled and processed using custom scripts and aligned against the IMGT germline references using IMGT/HighV-QUEST version v1.1.1 [12].

### Sample preparation, sequencing and processing of multiple sclerosis data

Samples from subjects M2, M3, M4, and M5 were collected from autopsy material that included central nervous system and draining cervical lymph node tissue derived from patients with multiple sclerosis [20]. Sequencing was performed as described in [20]. Briefly, V_H_ mRNA was selectively amplified by PCR using IGHV and IGHC region specific primers with 15 nucleotide unique molecular identifiers (UMIs). Amplicons where sequencing on the Illumina MiSeq platforming using the 2×250 kit according to the manufacturer’s recommendations. The version of the sequence data used here was previously used to generate lineage tree topologies as simulation constraints [21]. Briefly, sequence data was processed using pRESTO v0.3 [11] and Change-O v0.3.4 [17]. Reference alignment was performed using IMGT/HighV-QUEST v1.1.1 [12] with the February 4th, 2013 version of the IMGT gene database.

### Sample preparation, sequencing and processing of healthy monozygotic twin pair data

Subjects with identifiers beginning with TW represent five pairs of monozygotic twins whose BCR repertoires were previously sequenced from blood samples [29]. Peripheral blood was collected after obtaining written informed consent from all subjects, who participated in studies of licensed seasonal influenza vaccines under the Institutional Review Board approval at the Stanford University School of Medicine. Samples were prepared, sequenced and processed as described [29]. Briefly, FACS sorted cells were used to prepare sequencing libraries from RNA using a protocol employing 5’RACE and 10 nucleotide UMIs. Libraries were sequencing on the Illumina MiSeq platform using the 2×300 kit according to the manufacturer’s recommendations. UMIs and constant region primers were exacted from the raw reads using VDJPipe [30]. Further processing was performed using usearch [31], pRESTO [11], Change-O [17], and IMGT/HighV-QUEST v1.3.1 [12].

### Sample preparation, sequencing and processing of myasthenia gravis data and associated healthy controls

Subjects with identifiers beginning AR, MK and HD are from patients with myasthenia gravis with autoantibodies targeting the acetylcholine receptor (AR) or muscle specific kinase (MK) or from healthy controls (HD). Peripheral blood was obtained from subjects after acquiring informed consent and the study was approved by the Human Research Protection Program at Yale School of Medicine. Naive and memory B cells sorted from these subjects were previously published [22]. New data described here includes unsorted B cells from an additional subject MK06, and unsorted B cells from all subjects described in [22]. All samples were prepared, sequenced and processed as previously described [22]. Briefly, unsorted or FACS-sorted cells were used to prepare V_H_ and V_L_ sequencing libraries from mRNA using a protocol employing 5’RACE and 17 nucleotide UMIs. Libraries were sequencing on the Illumina MiSeq platform with the 2×300 kit according to the manufacturer’s recommendations, except for performing 325 cycles for read 1 and 275 cycles for read 2. Sequence data was processed using pRESTO v0.5.0 [11], Change-O v0.3.0 [17], SHazaM v0.1.2 [17], and IMGT/HighV-QUEST v1.4.0 [12] with the July 7, 2015 version of the IMGT gene database. Sequence data was deposited in the Sequence Read Archive (https://www.ncbi.nlm.nih.gov/sra) under BioProject accession PRJNA338795; sequencing runs used for this study are denoted A79HP, AAYFK, AAYHL, AB0RF and AB8KB.

### Genomic sequencing of predicted IGHV alleles

Genomic DNA was extracted using the Qiagen DNeasy Blood & Tissue Kit from the peripheral blood of subjects MK04, MK05 and MK06; peripheral blood was collected as part of the previously published myasthenia gravis study [22]. PCR primers were designed to fully amplify the exons and introns of each target IGHV gene locus (*IGHV1- 2*, *IGHV1-8*, *IGHV3-20* and *IGHV1-69*) from genomic DNA; sequences for each primer set are provided in Supplementary Table 2. PCR amplicons for each gene were generated individually from respective genomic DNA samples using the Qiagen HotStarTaq Kit (Cat. No. 203443), and subsequently cloned using the Invitrogen pCR4 TOPO TA kit (Cat. No. K457502). DNA was isolated from 4-15 clones per gene target, and sequenced from both ends using Sanger. Sequence chromatograms were viewed and analyzed using SeqMan Pro (DNASTAR 13.0.2).

### The updated TIgGER algorithm

The original TIgGER algorithm [8] was modified so that, for any set of sequences isolated from a single subject and best aligning to the same IgGRdb allele, the range of mutation counts analyzed would begin at the most frequent positive mutation count *m* and end at a mutation count of *m*+9. (If *m* = 1, the updated algorithm will behave as the original.) Additionally, any other mutation count at least 1/8 of the most frequent defines the start of a mutation range that is additionally analyzed, for improved sensitivity in cases where multiple novel alleles are assigned to the same IgGRdb allele; this mutation count may be either greater or less than the most frequent.

### Application of TIgGER to a human cohort

For novel allele detection and genotype inference, TIgGER was applied on functional, unique sequences with detectable junction sequences. For each sample, the “findNovelAlleles” function with default parameters was applied with IMGT IGHV germline reference (downloaded on May 17, 2018). Next, the set of putative novel alleles were used in genotype inference using the “inferGenotype” function with default parameters. Alleles that were included in the resulting genotype, but were not present in the IgGRdb, were considered novel alleles.

### Calculation of distant allele detection sensitivity

Pooled pre-vaccination sequences from subject PGP1 (i.e., samples taken at -8 days, -2 days, -1 hour relative to vaccination and sequenced on the 454 platform) were used. This dataset was chosen because it did not show significant clonal expansions in response to vaccination; did not have sequencing primers extending into the 5’ ends of sequences, as was the case in the multiple sclerosis and twin subjects, and had also been sequenced using the MiSeq platform, giving us the most confidence in the true set of alleles carried by the subject. For all sequences that best aligned to a particular IGHV germline allele, a number of positions *n* between IMGT-numbered positions 1 and 312 (inclusive) were modified (“mutated”) in the germline being used by the updated TIgGER algorithm. Mutations of a nucleotide to itself we not allowed, in order to ensure *n* differences between the starting germline and the resulting sequence. This was done 100 times for each *n* between 1 and 30, to simulate a situation in which the nearest IgGRdb was *n* polymorphisms away from the novel allele to be discovered, with each iteration using a separate random set of polymorphisms. The fraction of times the correct allele was detected by TIgGER for each value of *n* versus those detected at *n* = 0 (i.e., when TIgGER is allowed access to all IgGRdb alleles) was averaged across each germline sequence tested to determine the sensitivity as a function of *n*. For example, if for *n* = 15, 100/100 mutated variants led to the proper detection of the germline allele for 19 of 38 alleles, and in the remaining 19 alleles 90/100 mutated variants led to the proper detection of the germline allele in each case, then the sensitivity at *n* = 15 would be calculated as (19*100% + 19*90%) / 38 = 95%.

### Bayesian approach to genotyping

A Bayesian framework with a Dirichlet prior for the multinomial distribution was adapted to genotype inference. To model the possible allele distributions, up to four distinct alleles were allowed in an individual’s genotype (e.g., four alleles could correspond to a gene duplication with both loci being heterozygous). From the observed allelic frequencies, a posterior probability is calculated for a continuum of underlying biological models that set allelic distribution for each gene. For example, a gene can include two equally abundant alleles, or one allele that is twice abundant than the second one due to gene replication in one of the chromosomes (15). Prior distributions were initially set to reflect naive biological assumptions about the underlying dynamics that determine the allelic usage (see supplementary **Figure S1**). Following this initial approach, priors were modified by fitting empirically genotypes of the three subjects (all time points combined): PGP1, hu420143, and 420IV, constructed using the naive priors.

The posterior probability for each model is given by: 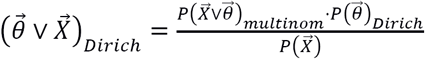, where 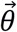 is the allele probability distribution and 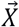 is the counts for the top four alleles. The certainty of the most probable model was calculated using a Bayes factor, 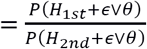 where H _1st_ and H_2nd_ correspond, to the most and second-most likely models, respectively. The larger the K, the greater the certainty in the model.

### Calculation of the Jaccard distance

To estimate distance between genotypes of two subjects a Jaccard distance was calculated in the following way: (i) for each gene, one minus the ratio between the number of shared alleles over the number of unique alleles from both samples was calculated. For example, for two genotypes with allele assignments *a* and *b* the Jaccard distance was defined as 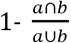. Genes that appeared in only one of the samples were excluded. (ii) The overall distance between two genotypes was calculated by a weighted average of all individual gene distances, where the weights are the mean of the two Bayes factors (K) for each.

## DISCUSSION

While the original TIgGER algorithm was very successful at detecting novel alleles, a significant limitation was that it could not detect novel V gene alleles that differed from known germline alleles by more than five SNPs. In addition, the original TIgGER genotyping approach was dependent on an arbitrary cutoff value for including genes in each subject’s genotype, and did not quantify the certainty of these genotype calls. Here we have described how modifying the “mutation window” in which the algorithm searches for mutation patterns that are indicative of polymorphisms was able to overcome the five mutations limitation. We also developed a Bayesian approach for genotyping that does not depend on a strict cutoff and provides a certainty level for each genotype call. We applied the updated algorithm to AIRR-seq data from 26 genetically distinct individuals [19, 20, 22, 29], and were able to identify 28 novel IGHV alleles. Although we showed on simulated data that TIgGER could detect alleles an arbitrary distance from known alleles, the most distant novel allele identified in this cohort contained three polymorphisms relative to the closest known IgGRdb allele. The fact that we did not discover more distant alleles is not necessarily surprising, as such sequences are expected to account for only ~10% of new alleles (using the distribution of distances between known IgGRdb as an estimate [8]). We expect that as TIgGER continues to be applied to datasets from more subjects, such alleles will be discovered.

The IMGT gene IgGRdb maintains its requirement of direct DNA-based allele evidence of any alleles to be included in the IgGRdb. We generated such validation for several TIgGER predictions, resulting in the inclusion of three novel IGHV gene alleles in IMGT: *IGHV1-2*06*, *IGHV3-20*03* and *IGHV1-69*17*. Validation of the other gene alleles discovered via AIRR-seq by TIgGER will be a priority going forward. While the IMGT standard for inclusion is intended to help ensure the quality of the IgGRdb, it inhibits the ability of the IgGRdb to benefit from the large number of non-IgGRdb alleles that are being rapidly discovered from AIRR-seq analyses. The Germline Gene Database (GLDB) Working Group of the AIRR Community is currently working to develop alternative criteria for judging the validity of Ig genes that are inferred from AIRR-seq data [18]. In the meantime, we have chosen to deposit the novel alleles we have detected into an alternative IgGRdb, the Immunoglobulin Polymorphism IgGRdb (IgPdb) [32]. Dependency on the completeness of IgGRdb can be reduced by TIgGER, as we demonstrated in deriving the majority of several subjects’ germline IGHV alleles starting from only a single gene allele per family. Further, a multiple alignment of the several sequences most-observed in a blood-based repertoire sample may be sufficient to remove the dependency on having a IgGRdb allele of each family, allowing for a more fully IgGRdb-blind derivation of alleles and V(D)J genotypes. Besides detecting several novel IGHV gene alleles in the genotypes of the 32 subjects in this study, we observed that no two IGHV genotypes appeared to be the same [33,34], barring those of the five pairs of monozygotic twins. It may be the case that IGHV genotypes alone are sufficient to uniquely identify a subject. This would additionally be improved if IGKV/IGLV genotypes, as well as D and J genotype were also determined; though theoretically possible, TIgGER has yet to be fully tested on J genes. However, we observed notable variation even in the genotypes of monozygotic twins due to the depth of sequencing. Though we adapted a Bayesian approach that presents an additional criterion for evaluating the certainty level of the genotype (based on the K value), in order to accurately differentiate samples coming from different individuals additional work is still required. One direction for further improvement of sample differentiation, was suggested recently by applying a Bayesian approach to haplotype inference [34]. We were able to accurately separate samples based on their genotypes from the subjects in the influenza time course, but these methods are affected by the sequencing depth. The influence of sequencing depth on the genotype call and its associated K value, was assessed on a single gene and is shown in supplementary **Figure S2**. It remains unclear how to adjust the Jaccard distance cutoff on the basis of sequencing depth, and we hope to explore this question and integrate dataset-tailored cutoffs into TIgGER’s genotyping functionality in the future.

Overall, we have expanded upon the capabilities of the TIgGER algorithm, demonstrated its persistent need in the analysis of AIRR-seq data, and hope that it will continue to be of use to the AIRR-seq community. New features and capabilities to TIgGER, such as the detection of insertion-deletion polymorphisms and the detection of novel J gene alleles, are being tested. The latest version of TIgGER is available for download as an R package from The Comprehensive R Archive Network (CRAN; http://cran.r-project.org) with additional documentation available at http://tigger.readthedocs.io. TIgGER is part of the Immcantation framework (http://immcantation.org), which provide a start-to-finish analytical ecosystem for high-throughput AIRR-seq data analysis, and is also available through the Immcantation Docker container builds at https://hub.docker.com/r/kleinstein/immcantation.

## SUPPLEMENTRY FIGURES

**Figure S1:**
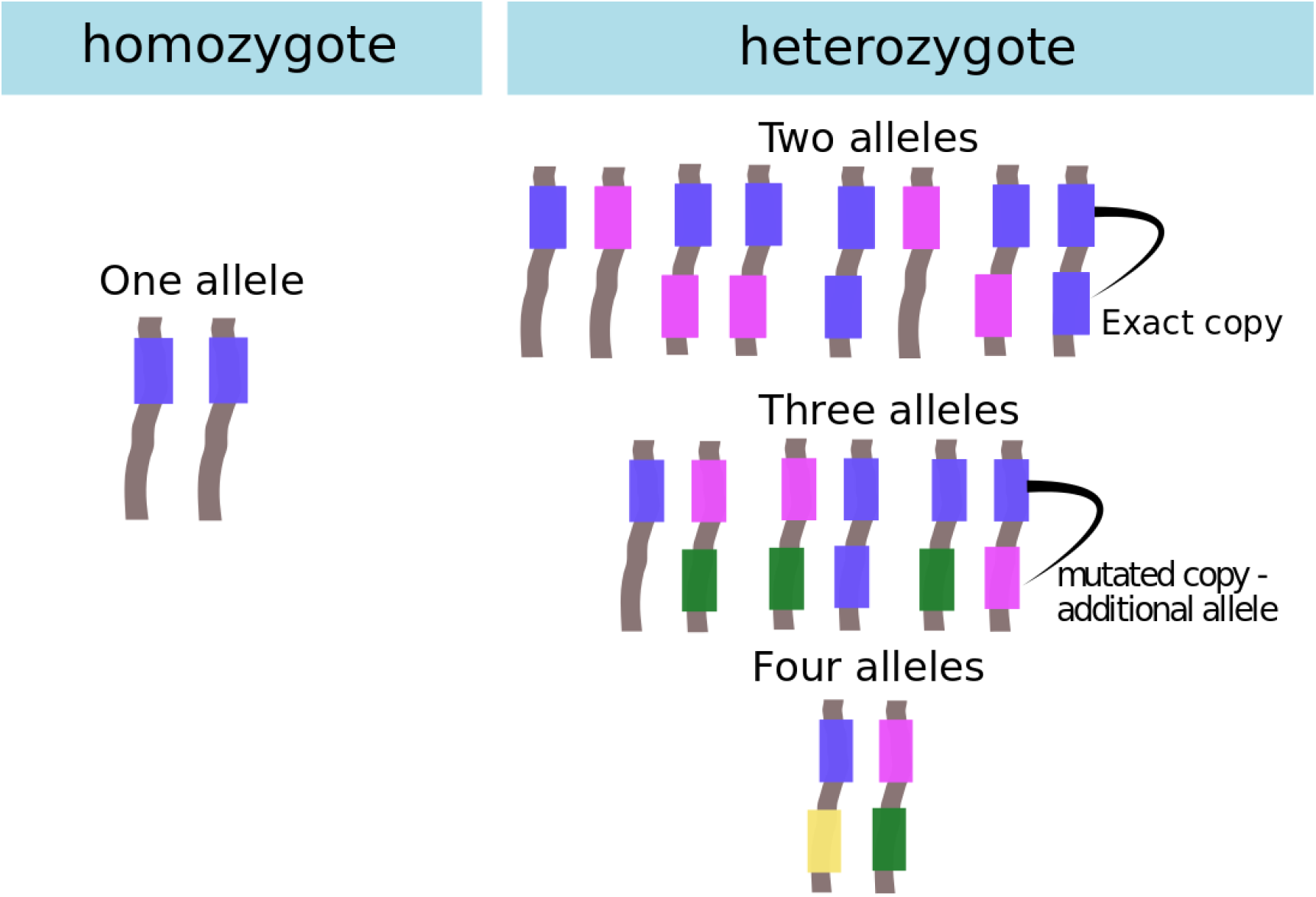
Models for genotype calls considered by the Bayesian method. The Bayesian genotyping method considers four potential models to explain the expression patterns of each gene. In the homozygote model (left), the same allele is carried on both chromosomes. In the heterozygote model (right), several possibilities are considered: in addition to two different alleles, one on each chromosome, up to two repeats of each allele on each chromosome were considered. Assuming polymorphisms can be present in each repeat, this results in up to four different alleles carried by a single individual.

**Figure S2:**
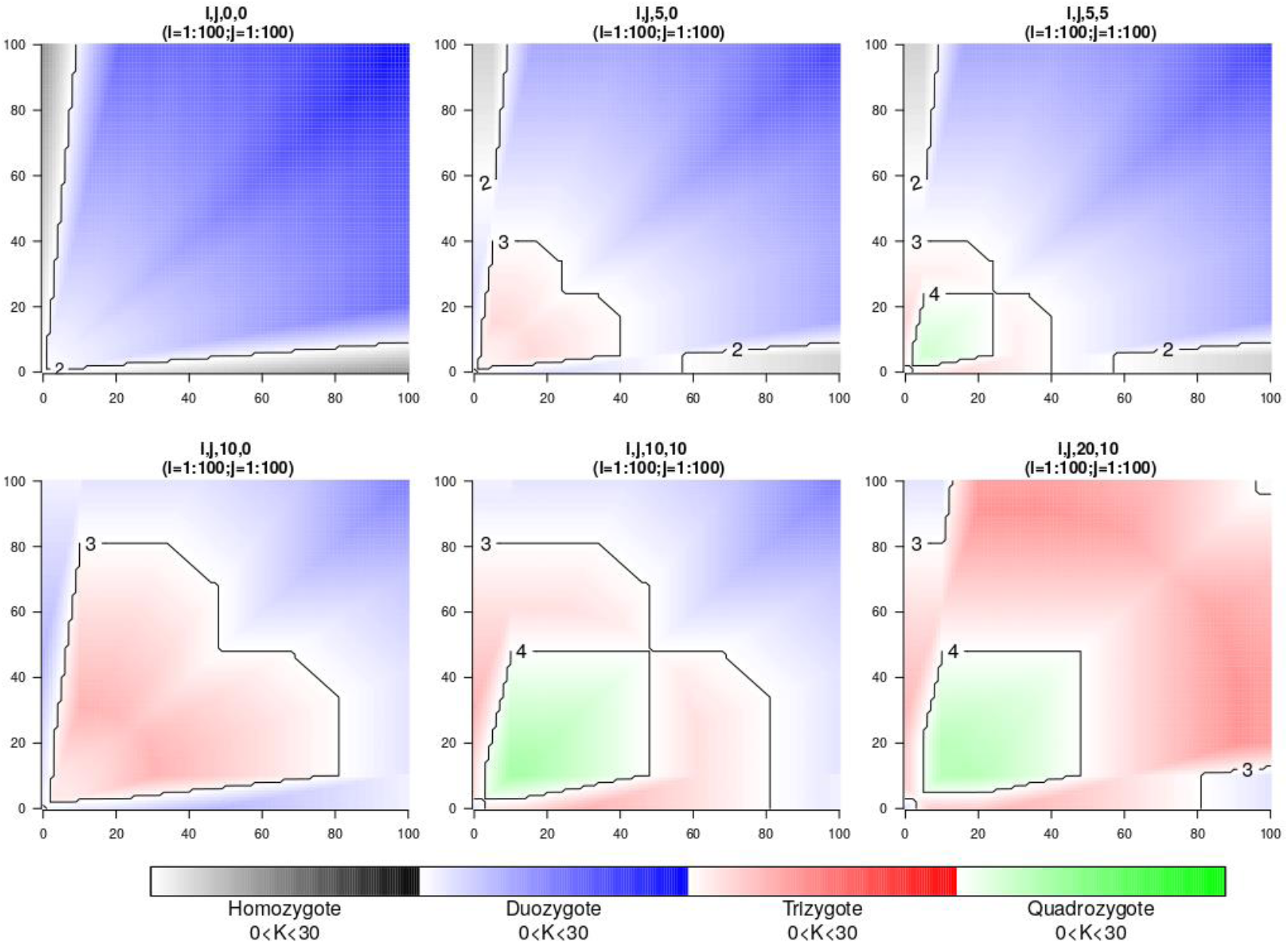
Genotype call and certainty levels as reflected by allele sequence counts for a single gene. The counts for the top two alleles in terms of frequency are represented by the Y and X axes, while counts for the third and fourth alleles are set for each panel and appear on the panel title. For example, for i,j,10,5 the counts of the third and fourth alleles are 10 and 5, respectively. The color reflects the genotype call: homozygote (black), douzygote - heterozygote with two alleles (blue), trizygote - heterozygote with three alleles (red) and quadrozygote - heterozygote with four different alleles (green). The intensity of the colors reflects the certainty level (K) of each genotype call.

## SUPPLEMENTRY TABLES

**Table S1:** Novel Immunoglobulin IGHV alleles discovered by TIgGER. For each biological sample in which a novel allele was discovered by TIgGER and determined to be in the genotype, the gene, nearest IgGRdb allele, and the polymorphisms that differentiate the novel allele from it are listed. (e.g., IGHV1-2 allele 02_T163C represents that the novel allele is most similar to IGHV1-2*02 and contains a T → C polymorphism at position 163.) The fourth and fifth columns indicate the number of unique sequences from the dataset that represent unmutated examples of the novel allele and the number of unique sequences from the dataset that represent unmutated examples of any allele of the given gene that was included in the genotype. The rightmost column indicates if the polymorphisms resulted in amino acid (AA) changes relative to the nearest IgGRdb allele, which is listed in the third column.

| Data Set | Gene | Allele(s) | Unmutated<br>Sequence<br>Count<br>(Allele) | Unmutated<br>Sequence<br>Count<br>(Gene) | AA change –<br>replacement(R)<br>/silent(S) |
| --- | --- | --- | --- | --- | --- |
| MK08 | IGHV1-18 | 01_A190G | 29 | 77 | R |
| AR05 | IGHV1-18 | 01_A196G | 276 | 566 | R |
| M2 | IGHV1-18 | 01_T111C | 732 | 1849 | S |
| M4 | IGHV1-2 | 02_T163C | 96 | 227 | R |
| M5 | IGHV1-2 | 02_T163C | 276 | 281 | R |
| hu420143 | IGHV1-2 | 02_T163C | 819 | 2028 | R |
| HD13 | IGHV1-2 | 02_T163C | 58 | 100 | R |
| AR05 | IGHV1-2 | 02_T163C | 312 | 682 | R |
| MK04 | IGHV1-2 | 02_T163C | 60 | 94 | R |
| TW071A | IGHV1-69 | 01_G54A | 65 | 106 | S |
| MK08 | IGHV1-69 | 04_G48A_A163G_A244G | 38 | 111 | S,R,R |
| M3 | IGHV1-69 | 06_C191T | 103 | 268 | R |
| MK06 | IGHV1-69 | 06_C191T | 94 | 468 | R |
| MK05 | IGHV1-69 | 12_G48A | 44 | 250 | S |
| PGP1 | IGHV1-8 | 02_G234T | 1548 | 3500 | S |
| MK05 | IGHV1-8 | 02_G234T | 91 | 91 | S |
| PGP1 | IGHV2-70 | 01_T164G | 221 | 350 | R |
| M2 | IGHV3-11 | 03_C300T | 420 | 719 | S |
| 420IV | IGHV3-11 | 03_T13G | 1625 | 2690 | R |
| PGP1 | IGHV3-20 | 01_C307T | 361 | 424 | R |
| MK05 | IGHV3-20 | 01_C307T | 61 | 61 | R |
| HD10 | IGHV3-30 | 18_C75G | 42 | 116 | S |
| 420IV | IGHV3-43 | 01_A112G_C222T_A286G | 175 | 408 | R,S,R |
| MK05 | IGHV3-48 | 01_A39C | 79 | 133 | S |
| M2 | IGHV3-64 | 05_A210C_G265C | 98 | 104 | S,R |
| MK05 | IGHV4-30-4 | 01_T120C | 33 | 33 | S |
| TW070B | IGHV4-34 | 01_A220G | 47 | 412 | R |
| TW071A | IGHV4-34 | 01_A220G | 78 | 910 | R |
| TW071B | IGHV4-34 | 01_A220G | 74 | 766 | R |
| TW071A | IGHV4-34 | 01_A220G_A225G | 58 | 910 | R,S |
| TW059A | IGHV4-39 | 01_A291G | 21 | 182 | S |
| TW071A | IGHV4-39 | 01_A291G | 12 | 111 | S |
| TW070B | IGHV4-39 | 07_A220G_A225G | 43 | 258 | R,S |
| TW064A | IGHV4-39 | 07_C288A | 31 | 81 | S |
| TW070A | IGHV4-39 | 07_C300T | 19 | 109 | S |
| TW070B | IGHV4-39 | 07_C300T | 26 | 258 | S |
| TW064A | IGHV4-59 | 01_T288C | 21 | 85 | S |
| TW064B | IGHV4-59 | 01_T288C | 20 | 76 | S |
| TW046A | IGHV4-59 | 01_T288C | 38 | 62 | S |
| TW059A | IGHV4-59 | 01_T288C | 28 | 67 | S |
| **TW059B** | IGHV4-59 | 01_T288C | 26 | 62 | S |
| **TW070A** | IGHV4-59 | 01_T288C | 34 | 74 | S |
| **TW070B** | IGHV4-59 | 01_T288C | 46 | 120 | S |
| **TW071A** | IGHV4-59 | 01_T288C | 13 | 57 | S |
| **TW059A** | IGHV4-59 | 08_A291G | 16 | 67 | S |
| **TW059B** | IGHV4-59 | 08_A291G | 9 | 62 | S |
| **TW070B** | IGHV4-59 | 08_A291G | 13 | 120 | S |
| **MK02** | IGHV4-61 | 02_A234G | 27 | 27 | R |

**Table S2:**
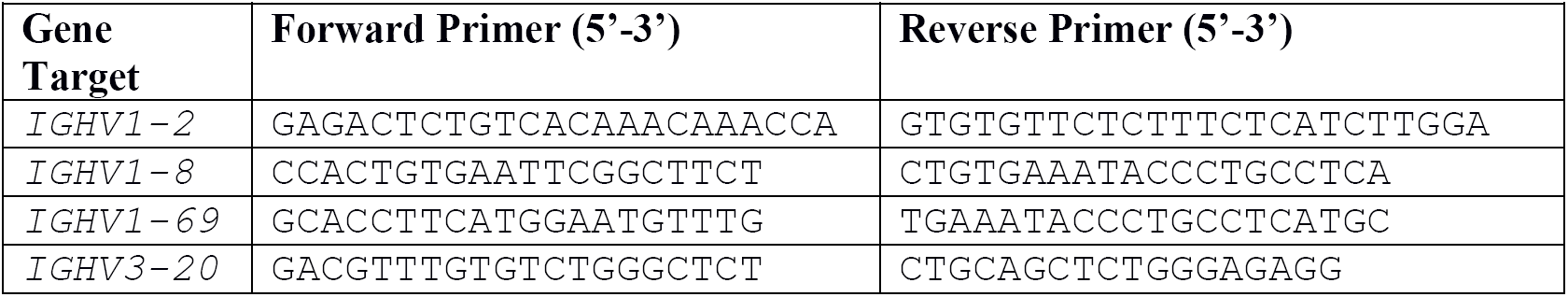
Primers used to fully amplify the exons and introns of each target IGHV gene locus.

